# A functional parcellation of the whole brain in individuals with autism spectrum disorder reveals atypical patterns of network organization

**DOI:** 10.1101/2023.12.15.571854

**Authors:** Andrew S. Persichetti, Jiayu Shao, Stephen J. Gotts, Alex Martin

**Affiliations:** Section on Cognitive Neuropsychology, Laboratory of Brain and Cognition, National Institute of Mental Health, National Institutes of Health, Bethesda, Maryland 20892

## Abstract

**BACKGROUND:** Researchers studying autism spectrum disorder (ASD) lack a comprehensive map of the functional network topography in the ASD brain. We used high-quality resting state functional MRI (rs-fMRI) connectivity data and a robust parcellation routine to provide a whole-brain map of functional networks in a group of seventy individuals with ASD and a group of seventy typically developing (TD) individuals.

**METHODS:** The rs-fMRI data were collected using an imaging sequence optimized to achieve high temporal signal-to-noise ratio (tSNR) across the whole-brain. We identified functional networks using a parcellation routine that intrinsically incorporates stability and replicability of the networks by keeping only network distinctions that agree across halves of the data over multiple random iterations in each group. The groups were tightly matched on tSNR, in-scanner motion, age, and IQ.

**RESULTS:** We compared the maps from each group and found that functional networks in the ASD group are atypical in three seemingly related ways: 1) whole-brain connectivity patterns are less stable across voxels within multiple functional networks, 2) the cerebellum, subcortex, and hippocampus show weaker differentiation of functional subnetworks, and 3) subcortical structures and the hippocampus are atypically integrated with the neocortex.

**CONCLUSIONS:** These results were statistically robust and suggest that patterns of network connectivity between the neocortex and the cerebellum, subcortical structures, and hippocampus are atypical in ASD individuals.

## Introduction

Autism spectrum disorder (ASD) is a developmental syndrome that affects a wide array of cognitive functions, ranging from core deficits in social-communication and restricted and repetitive behaviors to atypical sensory information processing ^1,2^. Researchers have had a lot of success using resting-state fMRI (rs-fMRI) functional connectivity methods to understand how such ASD-related deficits relate to atypical functional connectivity in specific brain networks. Resting-state functional connectivity data are especially well-suited to studying atypical neurophysiological dynamics in ASD because collecting it does not require researchers to design tasks to selectively probe the wide array of atypical cognitive functions and behaviors associated with ASD. In addition, resting-state functional connectivity data from multiple imaging sites can be aggregated and shared, as has been done in the large Autism Brain Imaging Data Exchange (ABIDE) data-sharing resource ^3,4^. However, despite impressive advances made using rs-fMRI to understand how ASD-related behavioral deficits correspond to functional networks in the brain, the field still lacks a whole-brain parcellation of functional networks in ASD individuals, thus leaving researchers to study brain connectivity in ASD by imposing network boundaries and regions of interest from brain maps that are based on data from typically developing (TD) individuals ^5–7^. A parcellation of the ASD brain is needed to provide a spatial map of whole-brain functional networks that is specific to the ASD group and can be used to make comparisons with parcellations of the TD brain.

Functional parcellations of human cortex have provided useful maps for studying the organization and function of the brain in TD individuals ^5–8^. It is well established that rs-fMRI activity is highly correlated within functional networks and these high correlations between regions within a network reflect direct or indirect anatomical connections ^9–15^. Thus, functional networks are identified and differentiated from one another in parcellation-based maps by grouping together brain regions that have similar patterns of rs-fMRI activity covariance with the whole brain. In this way, rs-fMRI parcellations reflect stable relationships between brain regions that can be used to map the functional organization of the human brain ^11,14,16–18^. In the current study, we provide an ASD-specific functional parcellation of the whole brain using rs-fMRI data.

Our parcellation uses high-quality rs-fMRI data from seventy ASD individuals that were collected using an imaging sequence specifically optimized to achieve high temporal signal-to-noise ratio (tSNR) across the whole-brain, including regions that usually suffer from relatively poor tSNR and distortions due to their close spatial proximity to the sinuses ^19,20^. We used a recently developed parcellation routine that intrinsically incorporates stability and replicability of the parcellation by keeping only network distinctions that agree across halves of the data over multiple random iterations ^21^. We also performed this parcellation routine on a control group of seventy TD individuals that were tightly matched to the ASD group in tSNR, in-scanner motion, age, and IQ, so that we could compare the functional networks identified in the ASD group with the results from the TD-group parcellation. After functionally parcellating the ASD and TD brains, we compared them on measures of network stability and differentiation of subnetworks.

## Methods

### Participants

Seventy individuals [mean (SD) age=19 (3.8) years; 14 female] who met the DSM-V criteria for ASD ^1^, as assessed by a trained clinician, were recruited for this experiment. In addition, seventy individuals with no history of psychiatric or neurological disorders [mean (SD) age=19.7 (3.7) years; 19 female] served as the TD control group. There were no significant differences between the two groups in age (t_(69)_=1.14, p=0.26) or overall IQ, as measured by the Wechsler Abbreviated Scale of Intelligence^22^. that was administered within one year of the scanning session to all participants [mean (SD), Full-score IQ, TD: 116.1 (11); ASD: 114.2 (12.9); t_(69)_=1.15, p=0.25]. Subsets of the resting-state data from these individuals have been used in a number of our previous studies ^21,23–27^. Informed assent and consent were obtained from all participants and/or their parent/guardian when appropriate in accordance with a National Institutes of Health (NIH) Institutional Review Board-approved protocol (10-M-0027, clinical trials number NCT01031407).

### MRI data acquisition and procedure

Scanning was completed on a General Electric Signa HDxt 3.0 T scanner (GE Healthcare) at the NIH Clinical Center NMR Research Facility. For each participant, T2*-weighted blood oxygen level-dependent (BOLD) images covering the whole brain were acquired using an 8-channel receive-only head coil and a gradient echo single-shot echo planar imaging sequence (repetition time = 3500 ms, echo time = 27 ms, flip angle = 90°, 42 axial contiguous interleaved slices per volume, 3.0-mm slice thickness, 128x128 acquisition matrix, single-voxel volume=1.7x1.7x3.0 mm, field of view = 22 cm). An acceleration factor of 2 (ASSET) was used to reduce gradient coil heating during the session. In addition to the functional images, a high-resolution T1-weighted anatomical image (magnetization-prepared rapid acquisition with gradient echo – MPRAGE) was obtained (124 axial slices, 1.2 mm^3^ single-voxel volume, 224x224 acquisition matrix, field of view = 24 cm).

During the resting scans, participants were instructed to relax and keep their eyes fixated on a central cross. Each resting scan lasted eight minutes and ten seconds for a total of 140 consecutive whole-brain volumes. Independent measures of cardiac and respiratory cycles were recorded during scanning for later artifact removal.

### fMRI data preprocessing

All data were preprocessed using the AFNI software package ^28^. First, the initial three TRs from each EPI scan were removed to allow for T1 equilibration. Next, 3dDespike was used to bound outlying time points in each voxel within 4 standard deviations of the time series mean and 3dTshift was used to adjust for slice acquisition time within each volume (to t=0). 3dvolreg was then used to align each volume of the scan series to the first retained volume of the scan. White matter and large ventricle masks were created from the aligned MPRAGE scan using Freesurfer ^29^. These masks were then resampled to EPI resolution, eroded by one voxel to prevent partial volume effects with gray matter voxels, and applied to the volume-registered data to generate white-matter and ventricle nuisance regressors prior to spatial blurring. Scans were then spatially blurred by a 6-mm Gaussian kernel (full width at half maximum) and divided by the mean of the voxelwise time series to yield units of percent signal change.

The data were denoised using the ANATICOR preprocessing approach ^30^. Nuisance regressors for each voxel included: six head-position parameter time series (three translation, three rotation), one average eroded ventricle time series, one “localized” eroded white matter time series (averaging the time series of all white matter voxels within a 15 mm-radius sphere), eight RETROICOR time series (four cardiac, four respiration) calculated from the cardiac and respiratory measures taken during the scan ^31^, and five Respiration Volume per Time (RVT) time series to minimize end-tidal CO_2_ effects from deep breaths ^32^. All regressors were detrended with a fourth order polynomial prior to denoising and the same detrending was applied during nuisance regression to the voxel time series. Finally, the residual time series were spatially transformed to standard anatomical space (Talairach-Tournoux).

To ensure that the fMRI data from both groups were high quality and matched, we measured the temporal signal-to-noise-ratio (tSNR) across the whole brain and a summary of in-scanner head motion using the @1dDiffMag program in AFNI. We calculated the tSNR in each voxel as the time series mean divided by time series standard deviation and selected participants from both groups that had high tSNR values across the whole brain. We used Diffmag (comparable to mean Framewise Displacement ^33^), which estimates the average of first differences in frame-to-frame motion across each scan run, to exclude participants with scores greater than 0.2 mm/TR. Both tSNR and in-scanner head motion were matched between the groups (Figure 1).

**Figure 1.**
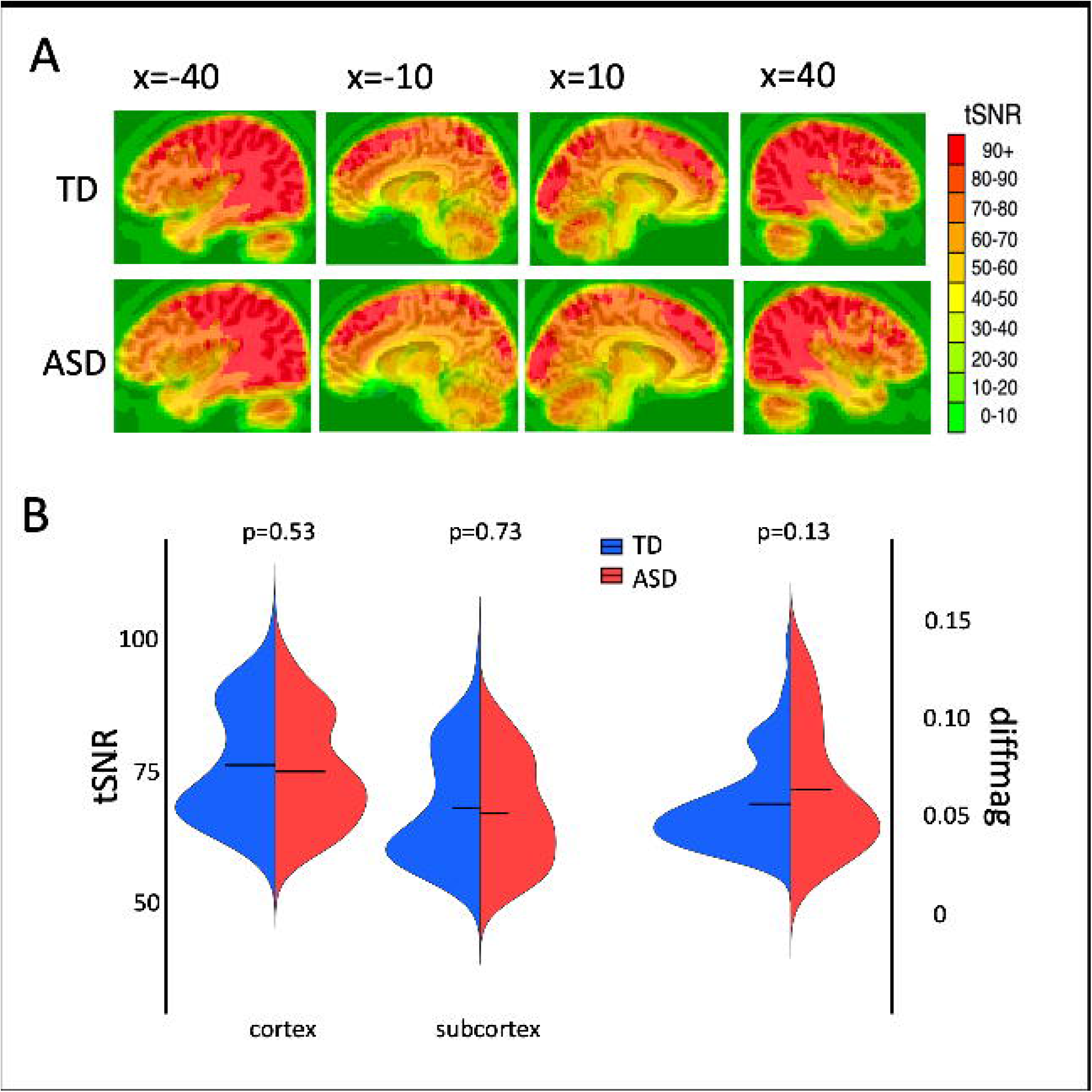
High-quality fMRI data were matched between the TD and ASD groups. **A)** Both groups had high temporal signal-to-noise ratio across the whole brain (tSNR – i.e., time series mean divided by time series SD). **B)** There were no significant differences in tSNR between the groups when averaged separately within the cortex and subcortex masks. Head motion was low in all participants and matched across groups (as measured using the DiffMag program in AFNI). Black horizontal lines in the violin plots represent the mean of each measure in each group.

### Resting-state parcellation routine

First, we made two masks using the Freesurfer cortical and subcortical segmentations ^34,35^: a cortical mask that included cerebellar voxels and a subcortical mask that included brain stem voxels (Figure 2A). Voxels with poor tSNR (< 10) and prominent blood vessel signal (identified from a standard deviation map of the volume-registered EPI data ^36^) were removed from the masks. The cortical mask was then downsampled to 6 mm^3^-resolution to speed up analysis run times, while the subcortical mask was downsampled to 3 mm^3^-resolution, because of its smaller starting volume.

**Figure 2.**
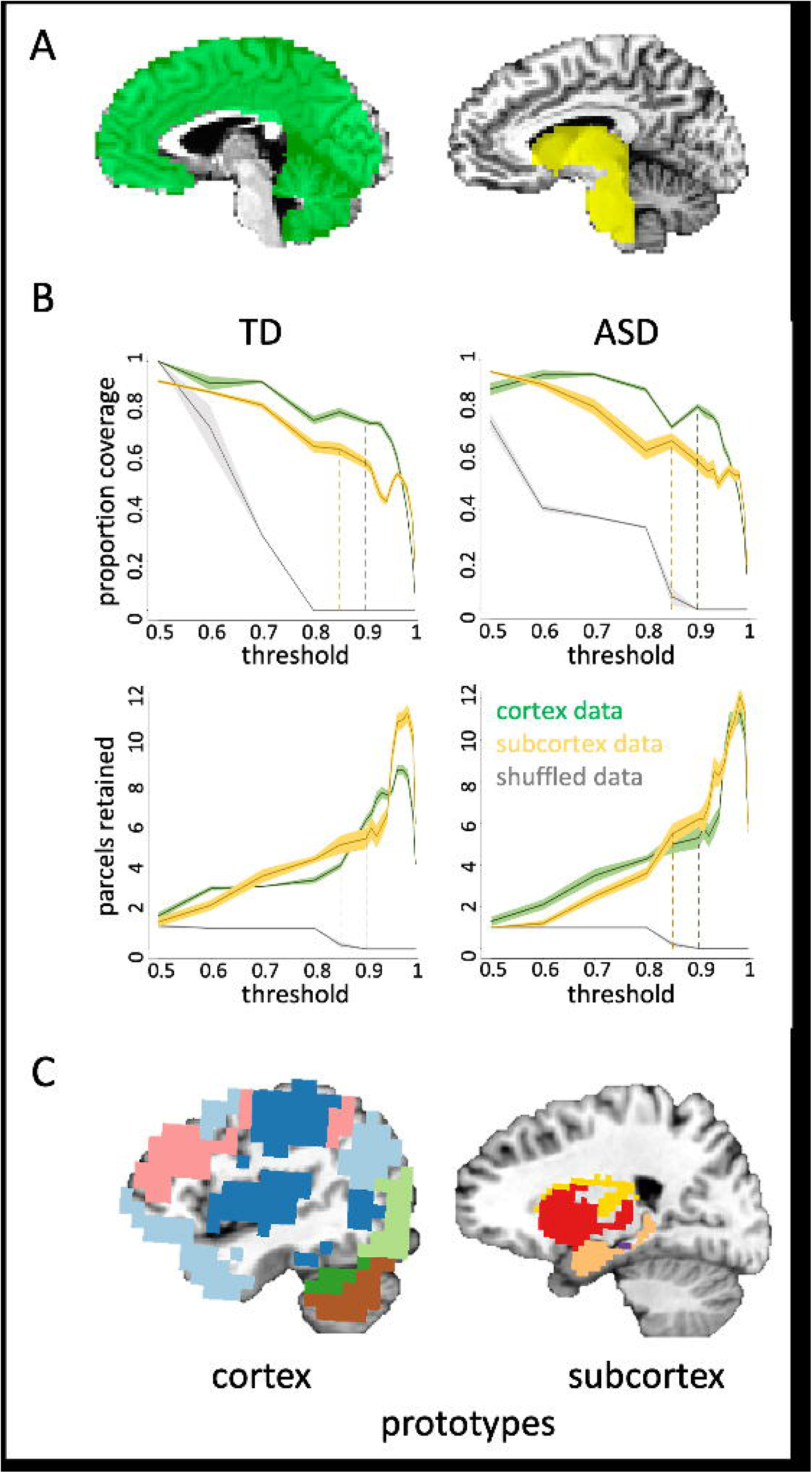
The initial parcellation routine. **A)** The parcellation focused on the cortical (left) and subcortical (right) masks separately. The cortical mask included all cortical and cerebellar voxels, while the subcortical mask included all voxels in the subcortex and brain stem. **B)** The spilt-half agreement curves were constructed across thresholds, picking the threshold that maximized proportion of coverage (i.e., number of voxels assigned a network prototype label) and the number of detected network prototypes (separately for cortical and subcortical masks). After ten iterations, one average parcellation of the retained network prototypes was formed, keeping any network that occurred in at least 50% of iterations. The proportion of coverage (top) and number of detected networks (bottom) were jointly optimized at the 90% threshold in the cortical mask and at the 85% threshold for the subcortical mask in both groups. The error in the line plots represents ±1 SEM. **C)** At this stage of the parcellation, we combined the masks so that all network prototypes were in the same space. This ensured that when we next ran the best-match procedure, so that every voxel in the whole brain was assigned a network label, any voxel could have a label that originated in either the cortical or subcortical mask.

We searched for functional network prototypes (i.e., sets of voxels in the group-averaged data with similar patterns of whole brain connectivity) across each mask using the InfoMap clustering algorithm ^37,38^. On each of ten iterations, the seventy participants per group were randomly split in half and group-average correlation matrices between the mask and whole-brain voxels were calculated for each half of data (done separately for the cortical and subcortical masks). These matrices were made square by correlating each column of the whole-brain x cortical (or subcortical) matrix with themselves. The real-valued correlation matrices were then thresholded into binary (0 or 1) undirected matrices at a range of threshold values (Figure 2B). The thresholded matrices of each half were then clustered using the InfoMap algorithm to form optimal two-level partitions (i.e., the optimal solution found over one hundred searches). A network prototype was counted as replicating across halves on each iteration if the Dice coefficient [Dice(x,y) = (2*(x∩y))/(x+y)] was ≥0.5, and the volume of the intersection was at least 2% of the size of the cortical or subcortical mask, respectively. The intersection of each network prototype that replicated across the two halves of data was retained for that iteration. After repeating the above steps for each of the ten iterations, one average parcellation of the retained network prototypes was formed, keeping voxels from any prototype that co-occurred in 50% or more of the iterations. Agreement curves were constructed across thresholds, and the threshold with the optimal proportion of coverage and number of detected prototypes was identified in each mask. We found that the split-half agreement and the number of detected prototypes were jointly optimized at the 90% threshold in the cortical mask and at the 85% threshold for the subcortical mask in both the TD and ASD group (Figure 2B). At this stage of the parcellation, every voxel is not guaranteed to have a network label due to the stringent requirements for replication across iterations described above. Thus, we next used a best-match criterion to ensure that all voxels were labelled in the end.

The detected network prototypes at the optimized thresholds in the cortical and subcortical masks were combined and then assigned to each voxel in the original 2 mm^3^ whole-brain mask using a best-match criterion. To do so, we first calculated the pattern of connectivity between each network prototype and the whole brain. The pattern of whole-brain functional connectivity for each network prototype was then compared with the pattern of connectivity from each voxel in the whole brain, and we assigned the label of the network prototype with the most similar pattern (Pearson correlation) to that voxel, provided the best match was within a threshold level of similarity (R^2^ > 0.5). Since the cortical and subcortical voxels were combined before assigning a final network label to each voxel, cortical voxels could, in principle, be labeled as belonging to a subcortical network, and vice versa, according to the best-match criterion.

### Calculating the Δ eta^2^ coefficient

We calculated the eta^2^ coefficient for every pair of voxels across the whole brain in each participant from both groups ^39^. The eta^2^ coefficient is defined as the ratio of variance in one variable that is explained by another variable. Thus, eta^2^ varies from 0 when there is no similarity between the variables and 1 when the variables are identical – i.e., a high eta^2^ indicates that two variables are similar to one another. We used eta^2^ to compare the whole-brain correlation maps between every pair of voxels and stored the eta^2^ coefficient in the first voxel location. If a pair of voxels are labelled *a* and *b*, then:

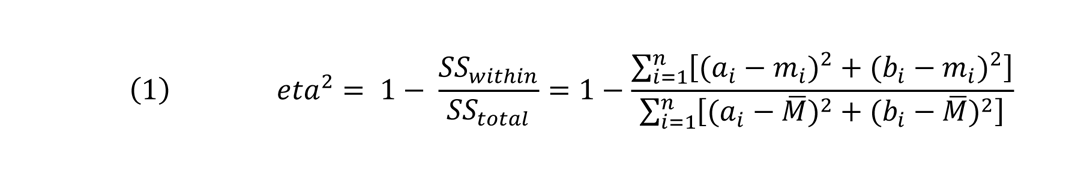

where *a_i_* and *b_i_* represent the position *i* in the correlation maps for *a* and *b*, respectively; *m_i_* is the mean value of the two maps at position *i*; and ^*m̅*^ is the grand mean value across the mean of the two correlation maps (i.e., *m)*. To calculate the Δ eta^2^ coefficient for each voxel, we averaged the eta^2^ coefficients between it and all other voxels from the same network (within-network eta^2^ coefficient) and separately averaged the eta^2^ coefficients between it and all other voxels from outside of the network (between-network eta^2^ coefficient). We then separately averaged the within-and between-network values across all voxels within a network and then subtracted the between-network eta^2^ coefficient from the within-network eta^2^ coefficient to get the Δ eta^2^ coefficient. Thus, if the parcellation identified meaningful functional boundaries in a group, then the Δ eta^2^ coefficient will be significantly positive. We calculated the Δ eta^2^ coefficient for each network in each participant. We then compared the average Δ eta^2^ coefficients between the ASD and TD groups.

### Calculating mean differences and null distributions to quantify group differences

To quantify the group differences in the number of networks found and the number of network-specific cortical voxels, respectively, we randomly split the data in half an additional one hundred times in each group and then compared the halves on each iteration. We did this separately in the cortical and subcortical masks. Doing so allowed us to compare one hundred observations from each group to obtain mean differences for our empirical observations between the groups. We also created null distributions by randomly labelling the two hundred halves of data either ASD or TD 25,000 times and generating a mean difference each time. We then evaluated the significance of our findings with permutation testing by comparing the real mean differences to the null distributions generated from the randomly shuffled split halves.

## Results

### Weaker differentiation of cerebellar networks in the ASD group

After detecting network prototypes in the subcortical and cortical masks separately (Figure 2C), we then combined the masks and found the best match to each prototype in every voxel across the whole brain in each group. In the TD group, we identified twelve whole-brain functional networks – six of the networks originated from prototypes in the cortical mask and the other six from prototypes in the subcortical mask (Figure 3A). By contrast, in the ASD group, we identified eleven whole-brain functional networks – five of the networks originated from prototypes in the cortical mask and the other six from prototypes in the subcortical mask (Figure 3B). The difference in the number of prototypes between the groups is due to the TD parcellation identifying two network prototypes in the cerebellum – roughly speaking, an anterior and posterior (Crus I/II and VIIB) prototype (dark green and brown, respectively, in Figures 3A&C) – while the ASD parcellation returned just one network prototype in the cerebellum (dark green voxels in Figure 3B). Consistent with our finding one less network prototype in the cortical mask of the ASD group compared to the TD group, but the same number of prototypes in the subcortical mask, the real mean difference in the cortex was greater than all 25,000 randomly shuffled mean differences in the null distribution (Figure 3D: mean difference=0.99, p<10^-5^), while the real mean difference in the subcortex did not differ significantly from the null distribution (Figure 3D: mean diff.=-0.01, p=0.76). Our finding of one less functional prototype in the cerebellum of the ASD group suggests that the cerebellar networks are weakly differentiated compared to the TD group. One consequence of this atypical differentiation of functional networks in the cerebellum of the ASD group is that an area of the posterior cerebellum is included in a cortical network that overlaps the default mode network (light blue in Figure 3B).

**Figure 3.**
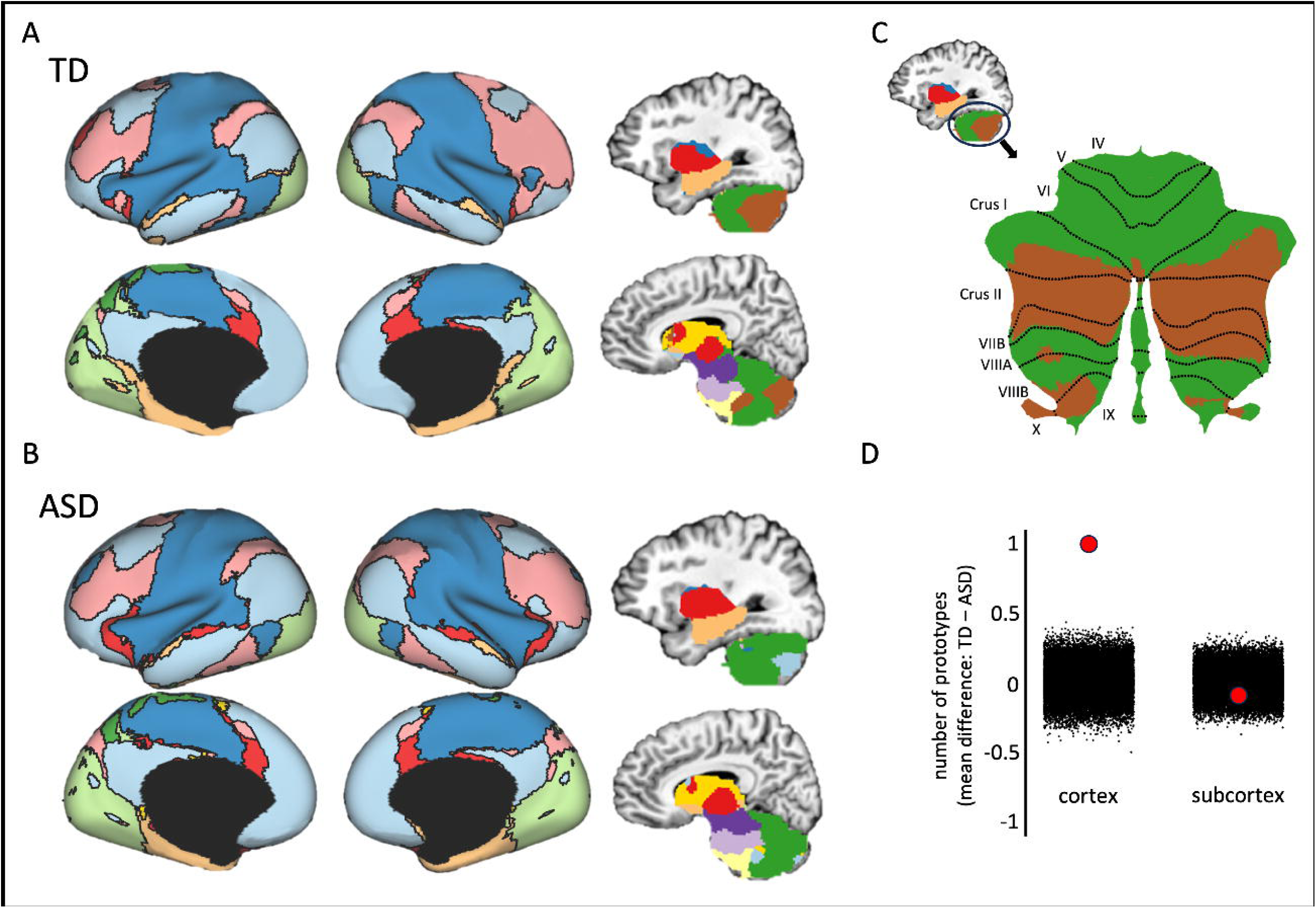
Resting-state parcellation of the whole brain in TD and ASD groups. **A)** Twelve networks were identified in the TD group parcellation – six originated from cortical prototypes and the other six from subcortical prototypes. Inflated brains were created using ther HCP Workbench (Marcus et al., 2011). **B)** Eleven networks were identified in the ASD group parcellation – five originated from cortical prototypes and the other six from subcortical prototypes. **C)** The cerebellar networks displayed on a flattened map of the cerebellum. The cerebellum was flattened using the SUIT toolbox (Diedrichsen 2006; Diedrichsen et al., 2009, 2011, 2015). **D)** The difference in the number of network prototypes between the groups was quantified by comparing the mean difference derived from one hundred random split halves of the data from each group (separately in cortical and subcortical masks) with a null distribution of 25,000 comparisons of the split-halves in which the group labels were randomly shuffled before obtaining the mean difference (black dots). The red dots are the actual mean difference in the number of prototypes between the groups. Positive values reflect a greater number of TD prototypes, while negative values correspond to more ASD prototypes.

Next, we examined three properties of the functional networks in both groups: 1) the degree of internal cohesion within each network, 2) the presence of differentiated subnetworks within each network, and 3) the spatial coverage of each network in the cortical and subcortical masks.

### Weaker network stability in the ASD group

We used the Δ eta^2^ coefficient as a measure of the degree to which the patterns of whole-brain connectivity are more similar for voxels within the same network compared to voxels from different networks. Thus, higher positive Δ eta^2^ coefficients reflect more cohesive and stable patterns of whole-brain connectivity across voxels from the same network. We found an overall significant decrease in the Δ eta^2^ coefficient in the ASD compared to TD group when averaging across all networks (independent samples t_(138)_=3.61, p<0.001, Cohen’s d=0.61) and this result was significant in both the cortical and subcortical masks when networks were averaged separately in each mask (both t’s>2.20, both p’s<0.05, both d’s>0.37). We next asked whether decreases of the Δ eta^2^ coefficient in the ASD group were significant in all functional networks or only a subset of networks. We found that the Δ eta^2^ coefficient was significantly lower in the ASD group within five functional networks (Figure 4: all t’s>2.48, p’s<0.01, d’s>0.42, FDR-corrected, q<0.05): hippocampal-cortical (pale orange), subcortico-cortical (red), sensorimotor (dark blue), fronto-parietal (pink), and anterior cerebellar (dark green). These results suggest that the whole-brain connectivity patterns from voxels within each of these networks, respectively, are less stable in the ASD compared to TD group. We confirmed this interpretation by showing that the reduced Δ eta^2^ coefficient in these regions of the ASD group are due to a greater decrease in within-network eta^2^ coefficients compared to between-network eta^2^ coefficients in the ASD group (Supp. Fig 1). Next, we tested whether this relative lack of network cohesion influences the organization of subnetworks within each of the affected large-scale networks.

**Figure 4.**
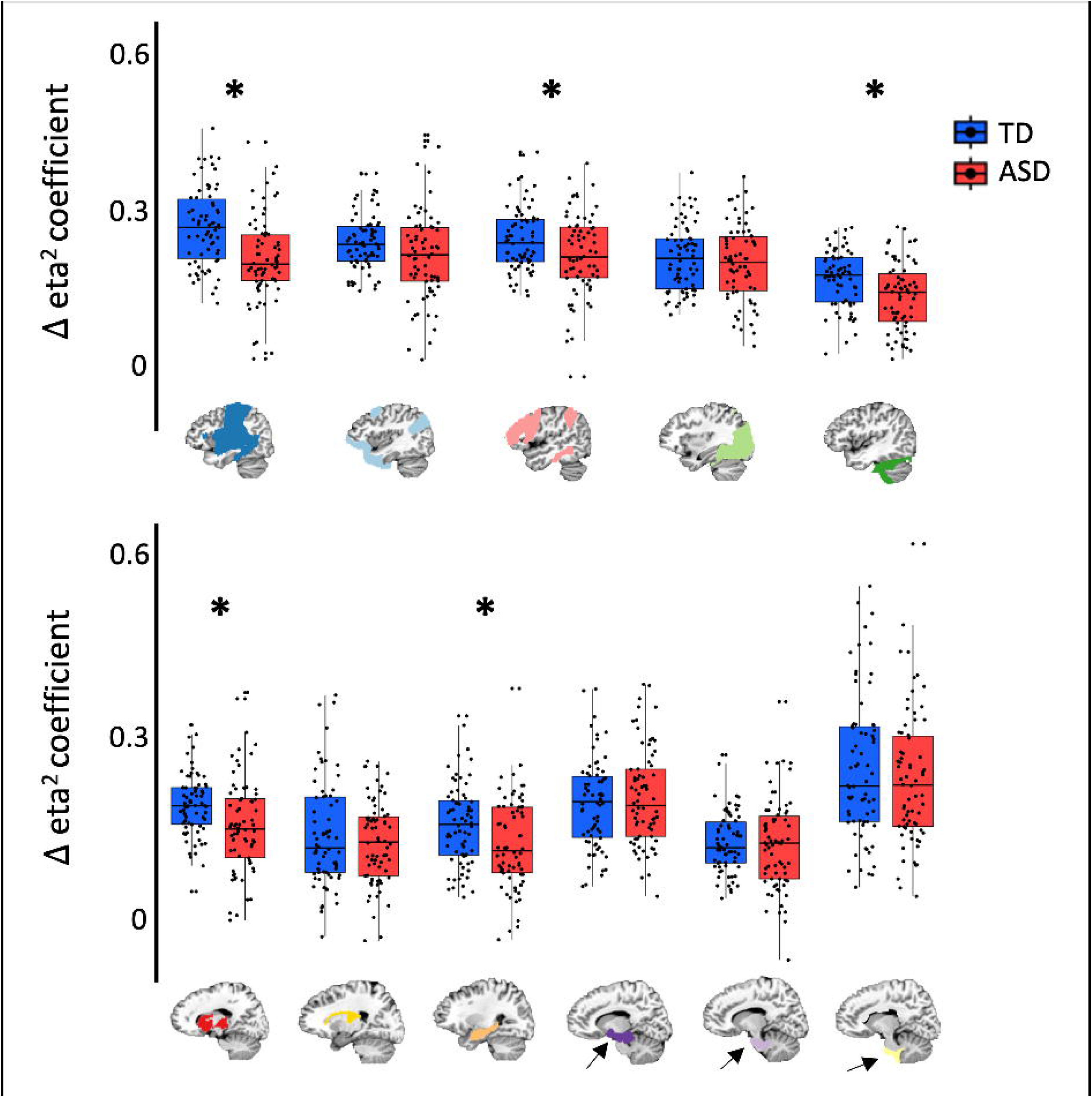
The Δ eta^2^ coefficients from each functional network in the TD and ASD groups. Asterisks represent a significant difference between the groups (p < 0.01). The networks that originated from cortical prototypes are on the top row and the networks that originated from subcortical prototypes are on the bottom row. In both rows, the networks are ordered by size (i.e., number of voxels). Note the sixth cortical network that corresponds to the posterior cerebellum is not shown because it is present in the TD group only.

### A relative lack of differentiated subnetworks in subcortex and hippocampus of the ASD group

To understand how weaker network cohesion in some large-scale networks of the ASD group might influence the organization of subnetworks within them, we used our parcellation method on each functional network in turn and evaluated the number of resultant subnetworks. Each large-scale network was treated as a mask and subjected to the same parcellation routine that was used to identify networks in the whole brain. We found differences in the number of subnetworks in the subcortical network that primarily overlaps the thalamus, putamen, and caudate nucleus (red mask in Figure 3) and the hippocampus (pale orange mask in Figure 3, but not in the sensorimotor and fronto-parietal networks (dark blue and pink, respectively, in Figure 3) that also showed a significantly lower Δ eta^2^ coefficient. The subcortex in the TD group divided the thalamus, putamen, and caudate nucleus into three subnetworks, while in the ASD group, the subcortex did not divide into subnetworks – i.e., it remained as one undifferentiated network (Fig. 5A). Similarly, the hippocampus in the ASD group was also less differentiated compared to the TD group (three vs. four subnetworks). Figure 5A shows that this difference in the number of subnetworks between the groups is most apparent on the long axis of the hippocampus. In both the subcortex and hippocampus, the mean difference in number of subnetworks between the groups (2 and 1, respectively) was greater than all 25,000 randomly shuffled mean differences in the null distribution (Figure 5B – both p’s<10^-5^). Note that we restricted our analysis space for these networks to the subcortex mask because, as will be demonstrated in the next section, each of these functional networks differ significantly in area of cortical coverage between the groups.

**Figure 5.**
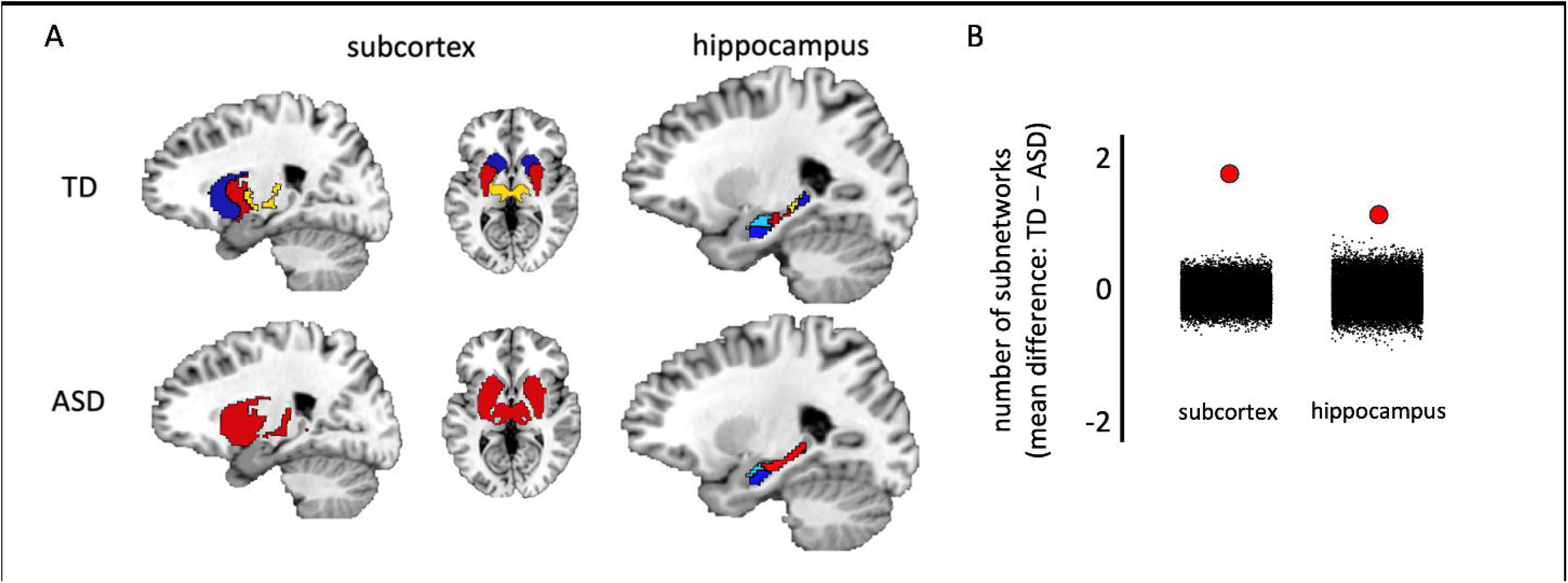
Differences in the number of subnetworks identified in the subcortex and hippocampus between the groups. **A)** As expected, the thalamus, putamen, and caudate nucleus were separated into three subnetworks in the TD group (top left), while the hippocampus was divided into four subnetworks along the long axis (top right). By contrast, the subcortex of the ASD group did not divide into subnetworks (i.e., it remained as one undifferentiated network – bottom left) and the hippocampus comprised one less subnetwork compared to the TD group (bottom right). **B)** The difference in the number of subnetworks between the groups was quantified by comparing the mean difference derived from one hundred random split halves of the data from each group (separately in the subcortex and hippocampus) with a null distribution of 25,000 comparisons of the split-halves in which the group labels were randomly shuffled before obtaining the mean difference (black dots). The red dots are the actual mean difference in the number of prototypes between the groups. Positive values reflect a greater number of TD subnetworks, while negative values correspond to more ASD subnetworks.

### Atypical subcortico-cortical and hippocampo-cortical integration in the ASD group

In addition, to weaker local differentiation of subnetworks in the subcortex and hippocampus of the ASD group, we next evaluated how the ASD and TD groups differed in the location and size of the neocortical areas that are integrated with the subcortical and hippocampal functional networks. To do so, we overlapped each network from the groups and calculated a ratio of voxels that intersected across groups versus voxels that were specific to one or the other group. We did this separately in the cortical and subcortical masks. For all networks with voxels in the subcortical mask, the ratio of intersecting to non-intersecting voxels was greater than half, so no further analyses were conducted on subcortical voxels. In the cortical mask, the ratio of intersecting to non-intersecting voxels was less than half only for the subcortical and hippocampal networks – i.e., these networks included more group-specific cortical voxels than voxels that intersect between the groups (Fig. 6A). For each network, the mean difference between the number of cortical voxels in the TD and ASD group was greater than all 25,000 randomly shuffled mean differences in the null distribution (Fig. 6B, mean differences, subcortical=1231.5 voxels, hippocampus=4974.6 voxels, both p’s<10^-5^). A map of the cortical voxels belonging to the subcortical network shows that there are more voxels that independently belong to the ASD group than belong to the TD group or intersect between the groups (Fig. 7A). These ASD-specific voxels are mostly located in and around the dorsolateral temporal cortex and the insula. By contrast, a map of the cortical voxels belonging to the hippocampal network shows that there are more voxels that independently belong to the TD group than belong to the ASD group or intersect between the groups (Fig. 7B). These TD-specific voxels are mostly located in lateral parieto-occipital cortex, around the retrosplenial cortex and parieto-occipital sulcus, and anterior lateral temporal cortex. These results show that the subcortical and hippocampal functional networks in individuals with ASD exhibit atypical connectivity patterns to the cortex.

**Figure 6.**
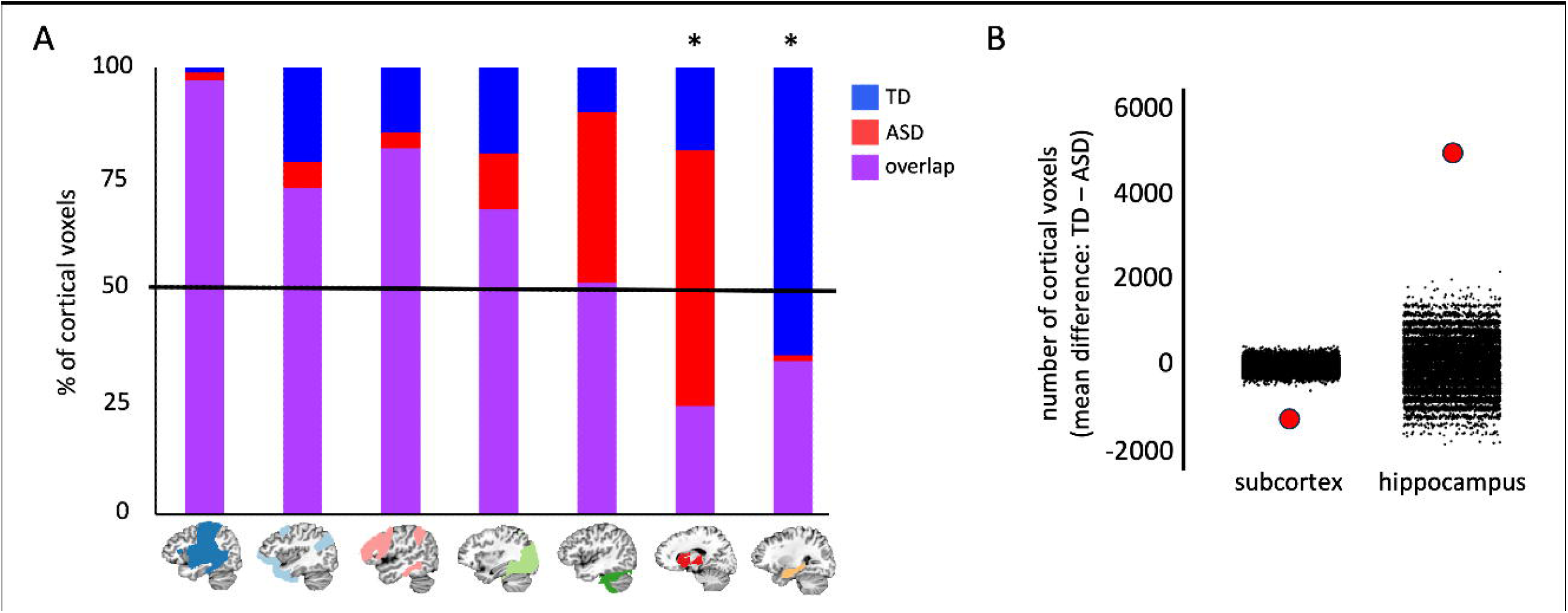
**A)** The percentage of intersecting and group-specific cortical voxels in each network. Of the eleven networks present in both groups (excluding the posterior cerebellar network not found in ASD), seven networks were present in the cortex, while four subcortical networks did not include more than fifty cortical voxels. All but two of the networks that were present in the cortex included more voxels that were intersecting between the groups than were exclusive to either group. By contrast, the network originating from a prototype primarily overlapping the thalamus, putamen, and caudate nucleus (red) included more cortical voxels that were exclusive to the ASD group, while the network originating from a hippocampal prototype included more cortical voxels that were exclusive to the TD group. **B)** The difference in the number of cortical voxels in the subcortical and hippocampal functional networks, respectively, between the groups was quantified by comparing the mean difference derived from one hundred random split halves of the data from each group (separately in the subcortex and hippocampus) with a null distribution of 25,000 comparisons of the split-halves in which the group labels were randomly shuffled before obtaining the mean difference (black dots). The red dots are the actual mean difference in the number of cortical voxels between the groups. Positive values reflect a greater number of TD subnetworks, while negative values correspond to more ASD subnetworks.

**Figure 7.**
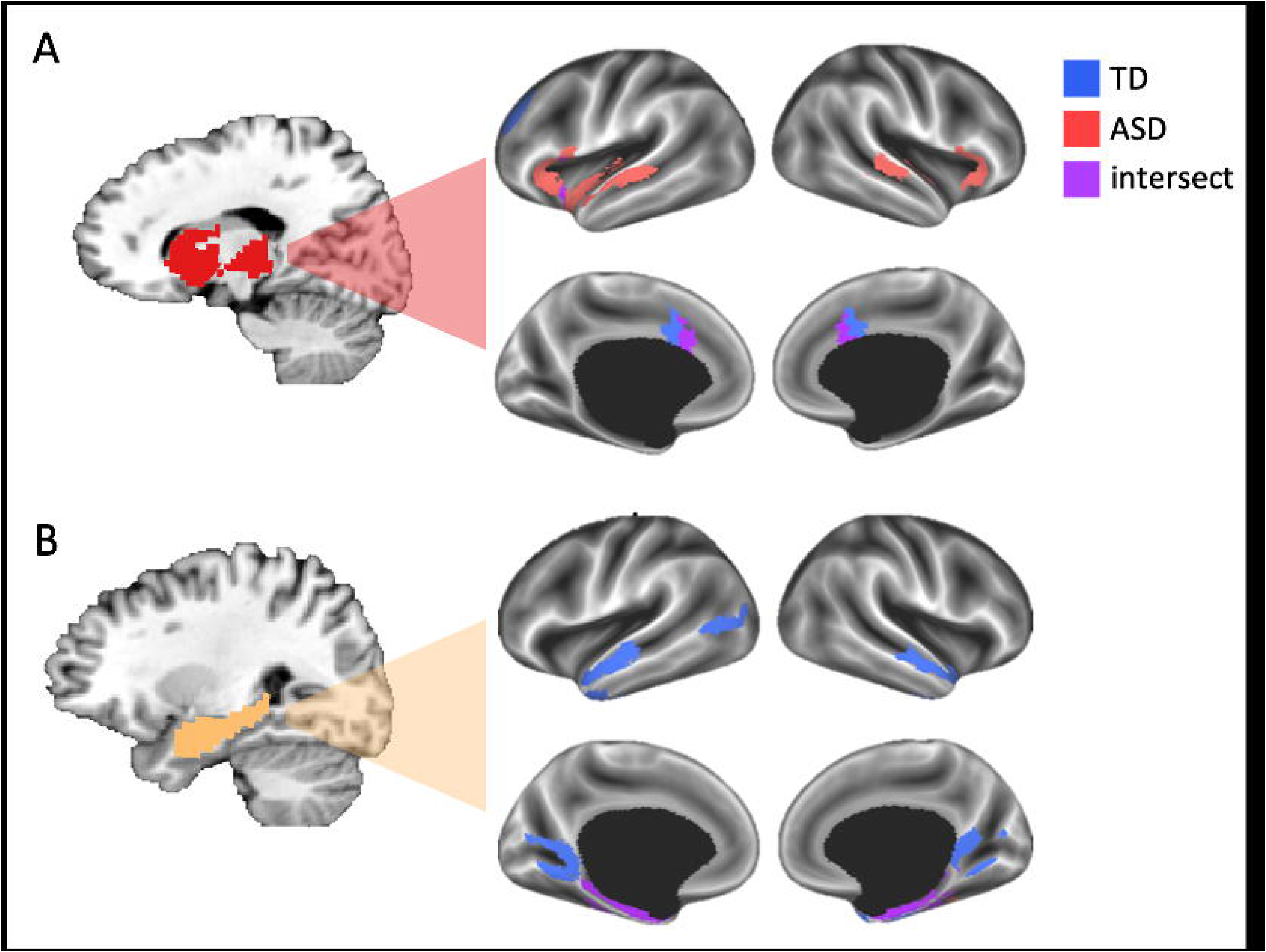
Group-specific cortical voxels in the subcortical and hippocampal networks. **A)** A network originating in the subcortex, primarily overlapping the thalamus, putamen, and caudate nucleus, is connected to more cortical voxels in the ASD than TD group. **B)** A network primarily overlapping the hippocampus, is connected to more cortical voxels in the TD than ASD group.

## Discussion

We used high quality rs-fMRI data and a robust parcellation routine to identify functional networks across the whole brain in individuals with ASD and tightly matched TD controls. We compared the functional networks from each group and focused on three atypical features of the ASD brain: 1) whole-brain connectivity patterns are less stable across voxels within select functional networks, 2) the cerebellum, subcortex, and hippocampus all show weaker differentiation of functional subnetworks, and 3) subcortical structures and the hippocampus are atypically integrated with the neocortex. These results were statistically robust and suggest that patterns of network connectivity between the neocortex and the cerebellum, subcortical structures, and hippocampus are atypical in ASD individuals.

The results mentioned above seem to be related in a straightforward way. Our finding of weaker cohesion within select networks of the ASD brain indicates that the patterns of whole brain connectivity from voxels across each of these networks are less stable compared to the TD group. Such instability is likely to be responsible for the relative lack of differentiation of subnetworks in the subcortical structures and hippocampus in the ASD group using our parcellation method. Interestingly, however, the lack of differentiation of the subcortical structures and hippocampus are coupled with opposite patterns of connectivity to the cortex – i.e., the subcortical structures are connected to more cortical voxels, while the hippocampus is connected to less cortical voxels compared to TD controls. The pattern of cortical connectivity from subcortical structures that is exclusive to the ASD group in our analysis overlaps with cortical regions that exhibited hyper-connectivity during rest and social tasks in prior reports – e.g., the insula and temporal lobes ^3,26,40,41^. The pattern of cortical connectivity from the hippocampus that is exclusive to the TD group in our analysis overlaps with cortical regions that exhibited hypo-connectivity between the hippocampus and cortex during episodic memory retrieval tasks ^42,43^. Intriguingly, some of the cortical regions missing from the hippocampal network in the ASD group in our analysis seem to overlap with scene-selective regions of cortex (i.e., retrosplenial and lateral occipitoparietal cortices ^44^), thus suggesting that this atypical network may be a neurobiological underpinning of reported behavioral deficits in scene construction and allocentric navigation in individuals with ASD ^45^. Overall, our results are more consistent with findings of atypical domain-specific network organization ^20,46–48^, rather than differences between the groups in global organizing principles, such as distance and strength of connectivity more generally ^49–51^.

In addition to the analyses presented here, the functional network map of the ASD brain can serve multiple functions in future studies. Since we demonstrated that the whole-brain functional network organization is significantly different between the ASD and TD groups, future studies can use our results to identify group-specific networks on which to focus their analyses, rather than combining data from the groups beforehand to identify networks common between them (as is typical of group ICA studies) or simply using parcels identified in TD groups in prior studies. The ASD-specific network map also provides a common spatial framework (or template) for integrating findings from studies that choose different regions (or networks) and/or behavioral deficits of interest. For example, several prior studies have reported unique patterns of behavioral correlates with each of the atypical networks that we focused on: The cerebellum – especially lobules Crus I/II and VIIB that we find undifferentiated in the ASD group – has been linked to deficits in social processing and communication ^52–55^. Subcortical structures – especially the thalamus – have been linked to deficits in social functions and sensory processing issues^56–58^. The hippocampus has been linked to deficits in episodic memory ^42,43,59^. For these reasons, the functional network maps of the ASD and TD brains from this study are freely available online (OSF: DOI 10.17605/OSF.IO/YT87Z).

## Acknowledgments

We thank Adrian Gilmore and Greg Wallace for insightful discussions and technical assistance. This work was supported by the NIMH Intramural Research Program. (#ZIA MH002920-09, clinical trials number NCT01031407). The authors declare no competing financial interests.

---

**Supp. Figure 1.**
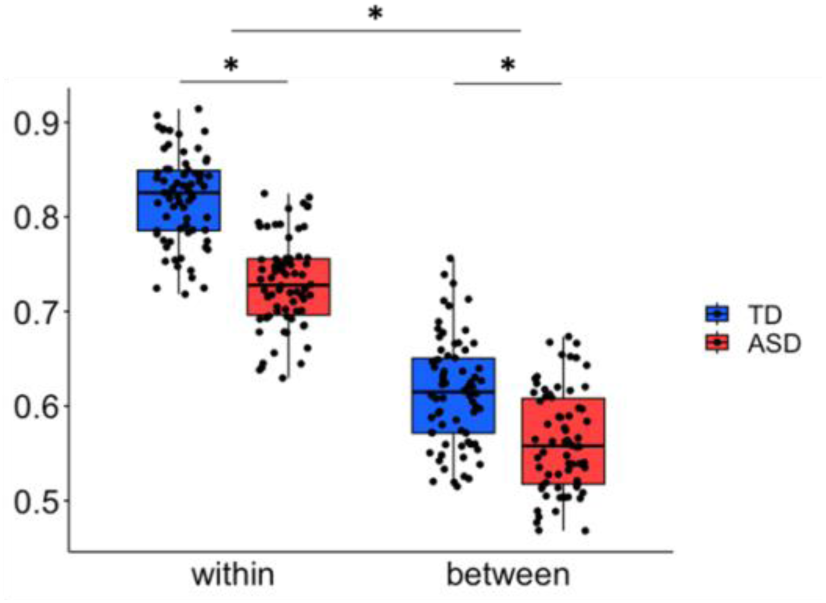
The Δ eta^2^ coefficient separated into within and between network voxels and then averaged across the five networks that showed a significant group difference in Figure 4. A significant Group x Within/Between interaction (p<0.0001) shows a greater decrease in within-network eta^2^ coefficients compared to between-network eta^2^ coefficients in these regions of the ASD group.

